# Coupling Mechanical and Metabolic Behavior in Huxley-type Muscle Models

**DOI:** 10.64898/2026.07.20.739506

**Authors:** Roeland T. Vos, Koen K. Lemaire, Luuk Vos, Arthur J. ‘Knoek’ van Soest, Dinant A. Kistemaker

## Abstract

Accurate prediction of both mechanical and metabolic behavior of muscle remains an important challenge in biomechanics. The typically employed Hill-type muscle models have shown limited success in this regard. Therefore, Huxley-type models, in which mechanical and metabolic behavior is linked through cross-bridge cycling, have gained renewed attention. Previous studies fitted the cross-bridge cycling rate parameter values of such models on mechanical behavior only. It seems reasonable to assume that accurate predictions of mechanical behavior in a Huxley-type model will also result in accurate predictions of metabolic behavior as both depend on the cross-bridge dynamics. Here, we show that this assumption does not hold. We simulated previously collected mechanical and metabolic data from an experiment where participants performed either isometric or dynamic knee extensions in the gravitational field. We modeled this experiment with a musculoskeletal model consisting of two segments driven by one Huxley-type muscle model. We obtained 10 sets of cross-bridge rate parameter values by systematically varying the value of one of the rate parameters and optimizing the values of the remaining rate parameters with respect to the mechanical behavior. We then compared the predicted mechanical and metabolic behavior between the 10 sets. The predicted mechanical behavior was similar for all 10 sets. However, the accuracy of the predicted metabolic behavior differed substantially between the 10 sets. Our findings illustrate that different sets of cross-bridge rate parameter values may lead to similar mechanical behavior. We conclude that this should be exploited to obtain accurate predictions of mechanical and metabolic behavior simultaneously in Huxley-type muscle models.

## Introduction

When studying the influence of metabolic energy consumption on human (loco)motion, musculoskeletal models can be a valuable addition to experimentation (Anderson & Pandy, 2001; Koelewijn et al., 2019; Miller, 2014; van den Bogert et al., 1998). Musculoskeletal models have been used to investigate hypotheses that would be difficult to test with a purely experimental approach, with studied tasks ranging from jumping (van Soest et al., 1993) to cycling (Neptune & van den Bogert, 1997) and level walking (Anderson & Pandy, 2001). Such studies typically use Hill-type muscle models (Hill, 1938) as they are computationally simple while providing accurate descriptions of the general mechanical behavior of muscles (Lemaire et al., 2016; Reuvers & Kistemaker, 2025a; Reuvers et al., 2025b; Reuvers et al., 2026; van den Bogert et al., 1998; Winters & Stark, 1987; Zajac, 1989). However, studying the metabolic energy expenditure of muscles using the Hill-type model is not straightforward: although Hill’s work was primarily focused on mechanical efficiency, the Hill-type model does not contain a direct relationship between the mechanical and metabolic behavior of muscles. Based on the heat terms in Hill’s original work (Hill, 1938), several extensions to the Hill-type model have been proposed to establish such a relationship (Bhargava et al., 2004; Lichtwark & Wilson, 2005; Uchida et al., 2016; Umberger et al., 2003; Umberger & Rubenson, 2011). Unfortunately, predicted metabolic energy expenditure for the same experimental task varies widely between these models (Afschrift et al., 2025; Koelewijn et al., 2019; Miller, 2014). The difficulty with these extended Hill-type models is that the heat terms do not have a direct relationship with the energy consuming processes in the muscle but are instead parameterized based on behavioral data obtained from dedicated experiments.

Huxley-type muscle models provide an alternative to the Hill-type model. In general terms, a Huxley-type model describes how muscles deliver force depending on the empirically observed cross-bridge attachment and detachment (Huxley, 1957). The advantage of the Huxley-type model is that it directly links the mechanical behavior of muscle to the metabolic energy expenditure of the cross-bridges (Huxley, 1957). Previous studies have also shown that predictions of the mechanical behavior made with Huxley-type models were accurate, and similar to those made with Hill-type models (Lemaire et al., 2016; van Soest et al., 2019). Furthermore, van Soest et al. (2019) showed that it is feasible to use the Huxley-type model in whole body musculoskeletal simulations of jumping, although the usage of Huxley-type models made these simulations computationally expensive. Building on the accurate prediction of mechanical behavior, Lemaire et al. (2025) investigated predictions of metabolic behavior in mouse fiber bundles made with Huxley-type models: they found that the relative error in the predicted metabolic energy, averaged over all mice, was 20.3% of the metabolic energy measured during the experimental trials. Recently, van der Zee et al. (2024) have successfully used Huxley-type muscle models to investigate the main rate-limiting processes in the dynamics of muscle force production. Based on these results, accurate prediction of mechanical and metabolic behavior in large scale musculoskeletal simulations using the Huxley-type model might be possible.

In the classic Huxley-type model, the attachment and detachment rates are functions of the cross-bridge bond length (Huxley, 1957). The distribution of attached cross-bridges determines both the mechanical and metabolic behavior of the cross-bridges (Huxley, 1957). In previous studies, the rate parameter values were optimized with the goal of obtaining accurate predictions of mechanical behavior in *in vivo* settings (van Soest et al., 2019; Vardy et al., 2012). This approach could also be used when both mechanical and metabolic behavior are of interest in an *in vivo* setting. If the representation of the relationship between the mechanical and metabolic behavior in the model (i.e. one cross-bridge cycle requires the consumption of one ATP (Huxley, 1957)) is valid, then one might expect that an accurate description of the mechanical behavior by definition leads to an accurate description of the metabolic behavior. However, this line of thought neglects the fact that the same mechanical behavior can be produced by different shapes of the rate functions (Huxley, 1957) or even different sets of parameter values for given shapes of the rate functions (Lemaire et al., 2025). As such, given the relation between cross-bridge detachment and ATP consumption, each of these sets must lead to different metabolic behavior. Lemaire et al. (2025) suggested that the redundancy in the rate parameter values with respect to the mechanical behavior can be exploited to improve the accuracy of the metabolic behavior of the model without compromising the quality of the predictions of the mechanical behavior.

In this study, we show that different sets of rate parameter values can lead to similar mechanical behavior yet result in markedly different metabolic behavior. Furthermore, we show that rate parameter values can be chosen to obtain accurate predictions of both mechanical and metabolic behavior simultaneously. We simulated previously published experimental data (van der Zee et al., 2019), in which participants performed knee extension contractions under both isometric and dynamic conditions. These simulations were performed using a forward simulation model driven by a single Huxley-type muscle–tendon complex model.

## Methods

We simulated both experimental conditions performed in the study by van der Zee et al. (2019). First, we chose a typical Hill-type muscle model (van Soest et al., 1993) for the baseline mechanical behavior of a quadriceps muscle. We then fixed the value of the first detachment rate parameter and, for each fixed value, optimized the remaining rate parameter values of the Huxley-type model to match the force–velocity relationship of the Hill-type model. This procedure yielded 10 sets of rate parameter values. We then used these 10 sets of rate parameter values to simulate the two experimental conditions. Lastly, we compared the predicted mechanical and metabolic behavior obtained with the 10 sets of rate parameter values. Metabolic predictions were based on a simplified version of the metabolic power model described by Lemaire et al. (2025).

### Experimental data

We simulated two experimental conditions that were part of a previous study by our group (van der Zee et al., 2019). In that experiment, participants were seated in an inclined chair and performed knee flexion/extension motions (see Fig. 1(A) in van der Zee et al. (2019)), while a weight of 2 kg was attached to each ankle. In both conditions, participants were given real-time feedback on their knee joint angle and were instructed to track a periodic knee joint angle template. In the first condition, subjects performed an isometric contraction where the target knee joint angle was constant, resulting in a constant torque. In the second condition, participants tracked a periodic knee joint angle trace that, when followed perfectly, would have required the same constant torque. The net mechanical work done was zero in both conditions, as the participant performed a periodic motion in the field of gravity. However, and importantly, the amount of positive mechanical fiber work was zero in the first condition but was not in the second condition. During both conditions, metabolic power was estimated using indirect calorimetry. For each condition, the net metabolic power was calculated by subtracting the metabolic power estimated during rest.

**Figure 1:**
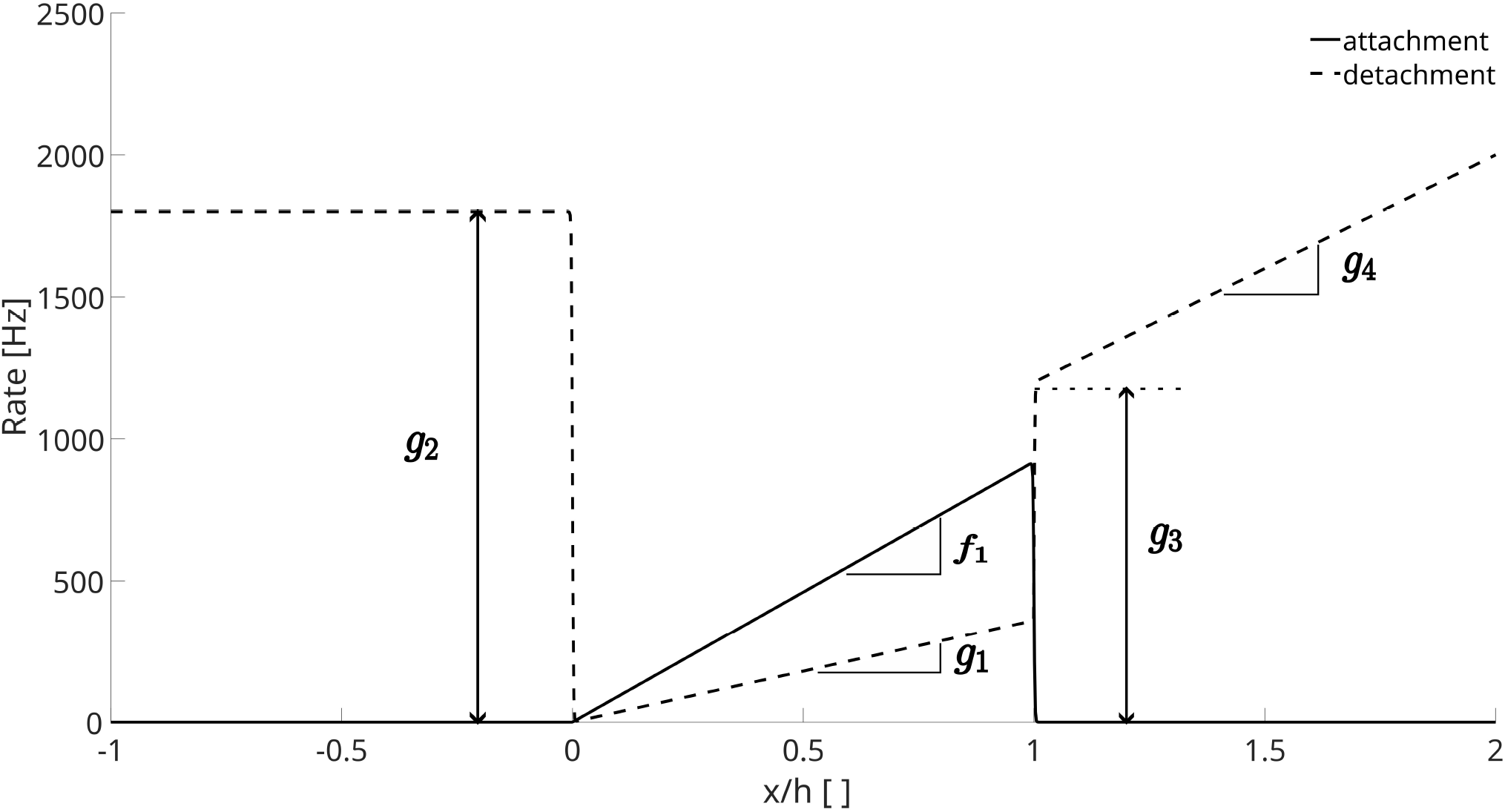
An illustration of the cross-bridge attachment and detachment rates as a function of normalized bond length, as determined by f_1_, g_1_, g_2_, g_3_ and g_4_.

### Simulation model

#### Overview

We devised a musculoskeletal model of the ‘average participant’ as described in (van der Zee et al., 2019). The skeletal part of the model consisted of two rigid segments connected through a friction-less joint. The first segment represented the upper legs, which were fixed. The second segment represented the lower legs, feet, and the added weights around the ankles. The model had one degree of freedom, which we described with the knee angle. The lower legs were driven by one Huxley-type muscle-tendon-complex (MTC) model representing the quadriceps and had normalized muscle stimulation as its only independent input. We excluded the antagonistic muscles from the model, as the measured electromyographical signal of these muscles was negligible in van der Zee et al. (2019). The Huxley-type MTC-model was similar to that used in Lemaire et al. (2025).

#### Attachment and detachment rates

The rate functions were chosen as described in Huxley (1957), with the only adjustment being the addition of two parameters (*g*_3_ and *g*_4_) to better fit the eccentric part of the steady-state force-velocity relationship, similar to Lemaire et al. (2025); see also Zahalak (1981). The rate functions used in this study were defined as follows:

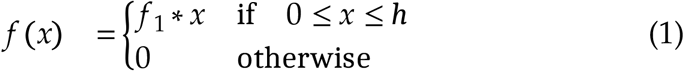

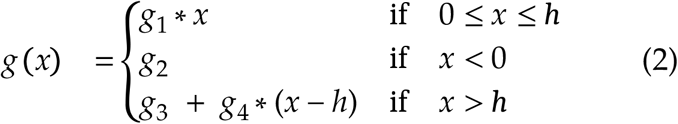

where *f x* and *g x* are the attachment and detachment rates as a function of cross-bridge bond length *x* and where *h* is the maximum attachment bond length (Huxley, 1957; Zahalak, 1981). The rate functions were slightly smoothed to reduce computational cost (see Lemaire et al. (2016)). An illustration of the rate functions is shown in Fig. 1.

The values of the rate parameters *f*_1_, *g*_2_, *g*_3_ and *g*_4_ were optimized to match the steady-state force-velocity relationship of the Huxley-type muscle to that of a Hill-type model of the same muscle (see the section *Numerical optimization procedures* for further details), similar to van Soest et al. (2019). The Hill-type model was implemented as done by van Soest et al. (1993). We searched for the value of *g*_1_ that led to the best match of predicted and experimentally measured mechanical and metabolic behavior using a grid search approach: for a range of values from 10 to 100 Hz with increments of 10 Hz, we performed the optimization procedures as described in the section *Numerical optimization procedures*. We chose to use *g*_1_ for the grid search as one of the simulated conditions consists of an isometric contraction. During an isometric contraction, the number of cross-bridge detachments, and therefore the amount of metabolic power needed for cross-bridge cycling, is almost exclusively determined by the value of *g*_1_. The sets of rate parameter values obtained from the optimization procedure will from this point on be referred to as *g*_1_*imposed*_*value*_, e.g. the set with *g*_1_ = 100 Hz is denoted as *g*_1_100_.

#### Metabolic power predictions

To predict metabolic power, we used a simplified version of the equation proposed by Lemaire et al. (2025); without these simplifications, the number of free parameters would be too large in comparison to the number of conditions. These simplifications did not have a substantial influence on our results (see Discussion). In the original version by Lemaire et al. (2025), two energy consuming processes were modeled: the pumping of calcium into the sarcoplasmic reticulum and the detachment of cross-bridges. The first simplification was to remove the term related to the pumping of calcium from the metabolic power equation. This was done because we expected the level of activation to be roughly equivalent between the isometric and dynamic conditions. We also did not account for mechanical detachments of cross-bridges at lengths greater than the maximal attachment bond length in contrast to Lemaire et al. (2025). Furthermore, we removed the scaling constant in the original equation of Lemaire et al. (2025) as we are only interested in the relative difference in metabolic power between the two conditions. The final equation for the metabolic power was thus as follows:

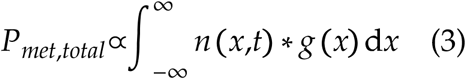

where *n* is the fraction of attached cross-bridges and 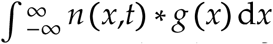 represents the total number of cross-bridges that detach per unit time. For each value of *g*_1_ considered (see above), we calculated the difference in *P*_*met,total*_ between the isometric and dynamic conditions, expressed as a percentage of the value of *P*_*met,total*_ during the isometric condition.

### Numerical optimization procedures

For each value of *g*_1_ considered, the values of the rate parameters *f*_1_, *g*_2_, *g*_3_ and*g*_4_ were optimized to match the steady-state force-velocity relationship of the contractile element in the Huxley-type model to that of the Hill-type model. The length of the contractile element was set to its optimum length (*lce*_*opt*_) and the active state (*q*), which represents the fraction of available cross-bridge binding sites, was set at 1. The cost function that was minimized was the sum of the squared differences in simulated steady-state CE-forces over a range of normalized contraction velocities ranging from -12 to 11 *lce*_*opt*_/s. In order to minimize the risk of arriving at a local minimum, for each value of *g*_1_, the optimization was performed with 20 different random initial guesses. The rate parameter sets with the lowest associated values of the cost function were then used for further analysis. The optimizations were performed using the simplex search algorithm of *fminsearch* in MATLAB R2024a.

Next, we investigated how well the mechanical behavior in the two experimental conditions of van der Zee et al. (2019) could be described for each of the values of *g*_1_ considered (see above). To that aim, we optimized the normalized muscle stimulation pattern (STIM) such that the sum of the squared differences between the predicted and experimentally observed knee extension torque was minimized. During the optimization of STIM, we imposed the length of the muscle-tendon-complex over time. As we are considering a periodic signal, we parameterized STIM (*t*) as a truncated Fourier series in the real form to simplify the optimization:

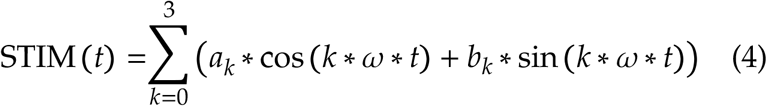

where ω equaled 2 ∗ π ∗ 0.5467 rad/s, the latter number being the movement angular frequency. The coefficients *a*_*k*_(_*k*_ = 0…3) and *b*_*k*_ (_*k*_ = 1…3) were the input variables for the optimization, which was performed using direct shooting and the SQP-algorithm of *fmincon* in MATLAB R2024a.

## Results

### Mechanical behavior

For all 10 values of *g*_1_, we were able to successfully optimize the values of the remaining rate parameters to closely match the steady-state force-velocity relationship of the Huxley-type MTC-model to that of a Hill-type model. Table 1 shows, for each of the values of *g*_1_ considered, the accompanying values of the rate parameters, as well as the corresponding RMSE of the simulated normalized forces.

**Table 1:** Sets of rate parameter values obtained from the numerical optimization procedure, as well as the corresponding values of the cost function to be minimized.

| $g_1$ [Hz] | $f_1$ [Hz] | $g_2$ [Hz] | $g_3$ [Hz] | $g_4$ [Hz] | RMSE [ ] |
| --- | --- | --- | --- | --- | --- |
| 10 | 1235 | 3165 | 683 | 1023 | 0.02604 |
| 20 | 1232 | 3154 | 686 | 1030 | 0.02603 |
| 30 | 1223 | 3155 | 684 | 1036 | 0.02603 |
| 40 | 1217 | 3148 | 688 | 1033 | 0.02607 |
| 50 | 1195 | 3159 | 679 | 1031 | 0.02608 |
| 60 | 1193 | 3146 | 685 | 1034 | 0.02610 |
| 70 | 1175 | 3169 | 682 | 1023 | 0.02608 |
| 80 | 1171 | 3151 | 684 | 1032 | 0.02607 |
| 90 | 1156 | 3160 | 682 | 1028 | 0.02608 |
| 100 | 1149 | 3158 | 680 | 1036 | 0.02606 |

Fig. 2 shows the simulated knee flexion angle (A), torque around the knee joint (B), and the relative free calcium concentration (C) during the dynamic condition for all 10 values of g_1_ considered. We can see in Fig. 2(B) that the simulated knee torque was close to the target torque for all 10 sets of rate parameter values, indicating that all 10 sets of rate parameter values led to similar mechanical behavior which was in line with our expectations.

**Figure 2:**
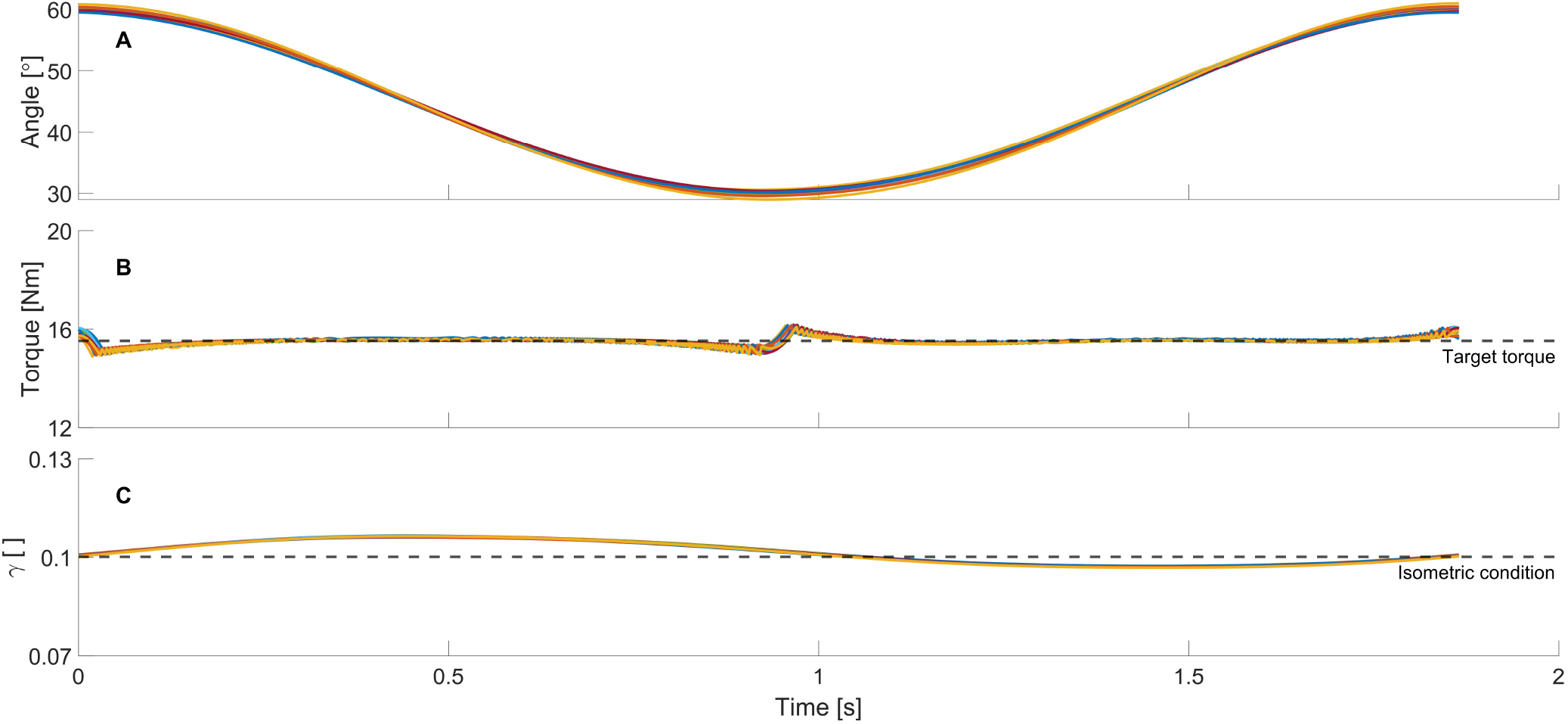
Simulation results for the dynamic condition for all 10 sets of rate parameter values (colored lines). **A:** The knee joint angle as a function of time. An angle of 0^∘^ indicates a fully extended knee. **B:** The knee extension torque as a function of time. The target value of the knee extension torque is shown as the dashed line. **C:** The normalized free calcium concentration as a function of time. The value of the simulated calcium concentration during the isometric condition is shown as the dashed line. Note that the mechanical behavior for all parameter sets was very similar, making the different lines hard to discern.

### Metabolic behavior

Fig. 3 shows, averaged over time, the fraction of attached cross-bridges (A) and the number of detachments (B) for two sets of rate parameter values (*g*_1_20_ and *g*_1_100_). Note that the integrals under the lines in Fig. 3(B) are proportional to the average metabolic power (eq. 3). For both sets of rate parameter values, we can see in Fig. 3(B) that the average number of detachments between *x*/*h* = 0 and *x*/*h* = 1 is similar between the isometric and dynamic condition. The increases in metabolic power during the dynamic condition therefore resulted from the additional detachments at *x*/*h* < 0 and *x*/*h* > 1. At *x*/*h* < 0 and *x*/*h* > 1, the number of detachments was slightly higher for *g*_1_20_ compared to *g*_1_100_. This increase in the number of detachments resulted from a larger fraction of attached cross-bridges when setting *g*_1_ = 20 (see Fig. 3(A)). Furthermore, the number of detachments in the isometric condition is lower for *g*_1_20_ between *x*/*h* = 0 and *x*/*h* = 1 compared to *g*_1_100_. Thus, compared to *g*_1_100_, the additional detachments at *x*/*h* < 0 and *x*/*h* > 1 had a relatively larger influence on the difference in metabolic power between the conditions. These findings explain why the predicted difference in metabolic power was higher for *g*_1_20_ compared to *g*_1_100_ (see Fig. 4).

**Figure 3:**
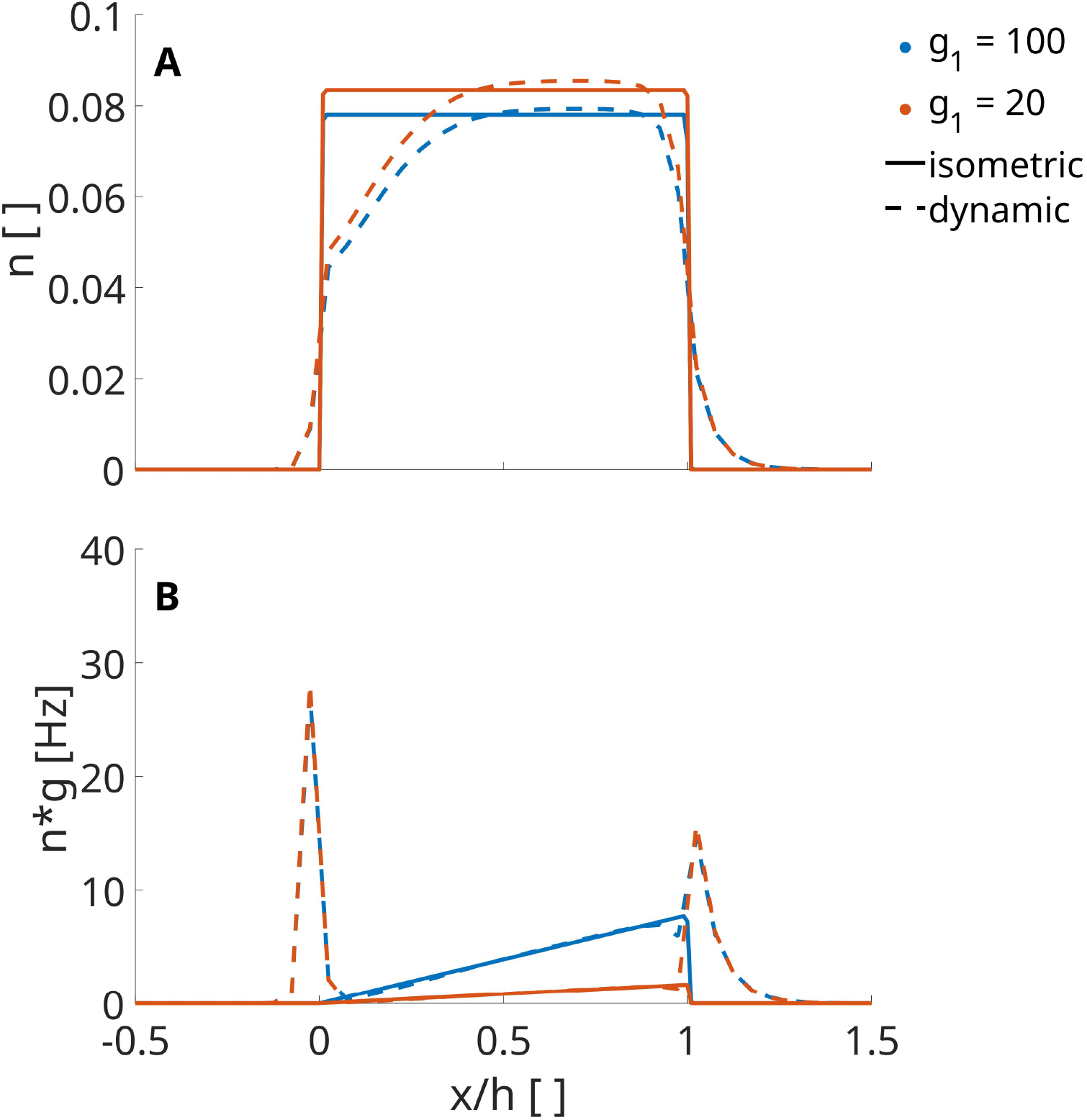
The fraction of attached cross-bridges (**A**) and number of detachments per unit time (**B**), averaged over time, as a function of cross-bridge bond length for two sets of rate parameter values (g_1_20_ and g_1_100_).

**Figure 4:**
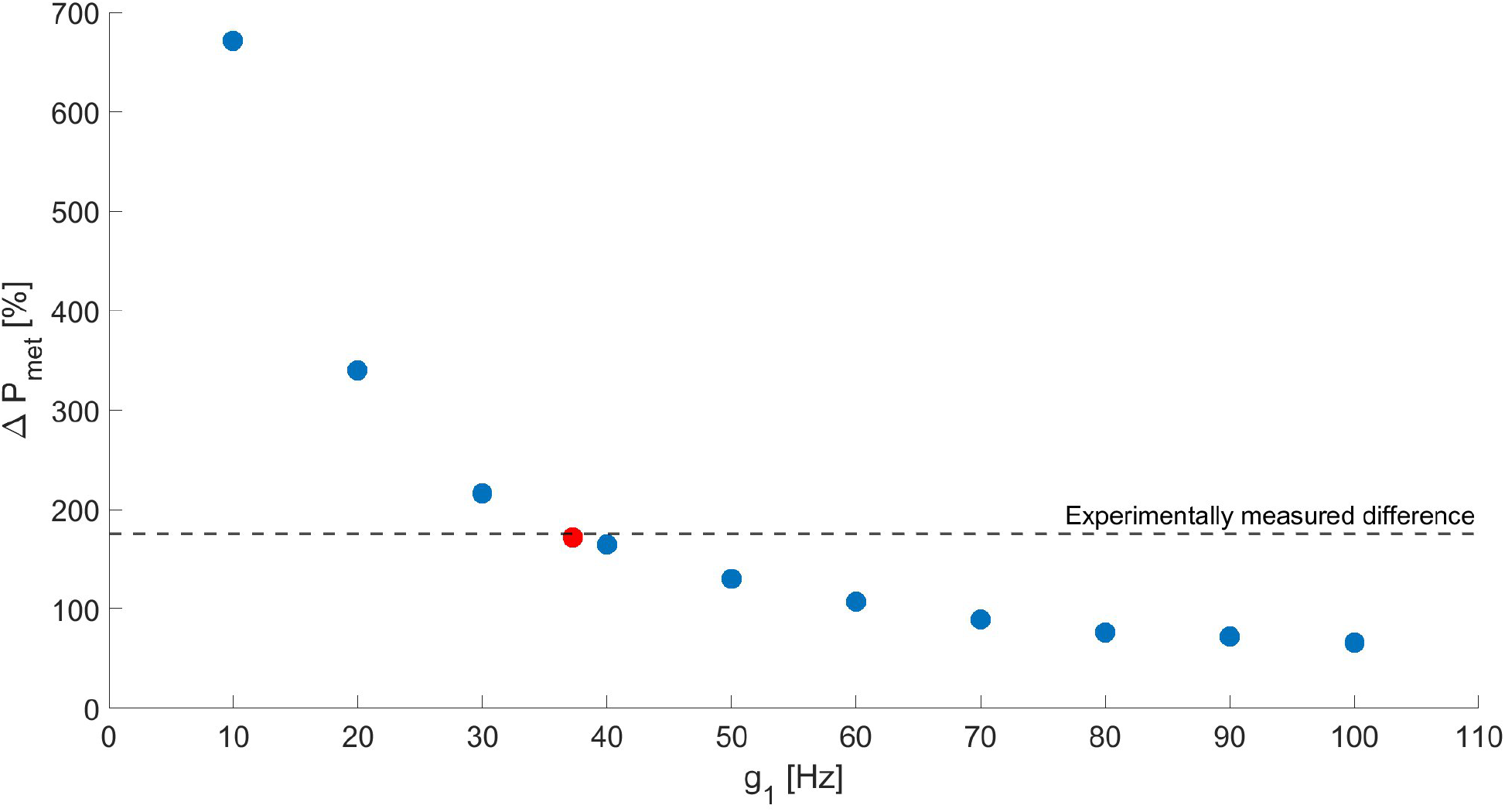
Relative difference in metabolic power between the isometric and dynamic condition as a function of the fixed rate parameter g_1_. The relative difference is expressed as a percentage of the predicted metabolic power in the isometric condition. The dashed line indicates the experimentally measured difference in net metabolic power between the conditions (van der Zee et al., 2019). The red dot indicates the predicted difference in metabolic power for the estimated optimal value of g_1_.

Fig. 4 shows, expressed as a percentage of the predicted metabolic power in the isometric condition, the predicted difference in metabolic power between the isometric and dynamic condition as a function of *g*_1_. For *g*_1_10_, the predicted difference in metabolic power was 671% of the metabolic power in the isometric condition. On the other hand, *g*_1_100_ resulted in a relative difference in metabolic power that was substantially lower, around 66%. *g*_1_40_ led to the best match of the predicted and experimentally measured difference in metabolic power between the conditions (Fig. 4): the predicted difference was 164%, while the experimentally measured difference in net metabolic power was 176% (van der Zee et al., 2019). As is obvious from Fig. 4, the match between the predicted and experimentally measured difference in metabolic power can be improved by further fine-tuning the value of *g*_1_. To illustrate this, we used piece-wise cubic spline interpolation to estimate the value of *g*_1_ that would lead to a perfect match between the predicted and experimentally measured difference in metabolic power between the conditions. Then, we performed the procedure as described in the Methods to obtain the predicted difference in metabolic power using this optimal value of *g*_1_. The estimated optimal value of *g*_1_ was 37 Hz, and the predicted difference in metabolic power was indeed very close to the experimentally measured difference, with the predicted value being 172% (see also the red dot in Fig. 4).

To summarize, while the 10 *g*_1_-imposed sets of rate parameter values resulted in very similar mechanical behavior, they resulted in rather different metabolic behavior.

## Discussion

### Overview

In this study, we aimed to show that the same mechanical behavior can be obtained using different sets of rate parameter values in a Huxley-type model. More importantly, we aimed to show that this can be exploited to obtain accurate predictions of both mechanical and metabolic behavior simultaneously. To do so, we first obtained 10 different sets of rate parameter values by fixing the value of *g*_1_ (the first detachment rate parameter, see Fig. 1), after which the values of the remaining rate parameters were obtained through numerical optimization with respect to the steady state force-velocity relationship. Then, we compared the predicted mechanical and metabolic behavior obtained with these 10 sets of rate parameter values to experimental data of van der Zee et al. (2019). We found that mechanical behavior was very similar for each of the 10 sets of rate parameter values, while the predicted metabolic behavior differed substantially between the 10 sets of rate parameter values. These findings indeed show that accurate prediction of mechanical behavior by a Huxley-type muscle model does not imply that the predicted metabolic behavior will be accurate as well. We will now first discuss the potential influence on our results of two choices that we made relating to simplification of the metabolic power model. Then, we will discuss other possible formulations of the rate functions. Lastly, we will discuss the implications of our findings with respect to the implementation of Huxley-type models for predicting mechanical and metabolic behavior of muscles in musculoskeletal simulations.

### Influence of model simplifications on the results

In the Methods, we explained that we used a simplified version of the model for metabolic power described by Lemaire et al. (2025). The two major simplifications we made were the assumption that cross-bridges would not mechanically detach at high bond lengths (i.e. they still require ATP to detach) and the exclusion of metabolic power consumption due to muscle activation. The inclusion of mechanical detachments would lead to a decrease in the optimal value of *g*_1_ with respect to the predicted difference in metabolic power between the two conditions. However, the aim of this study was not to pinpoint the optimal value of *g*_1_, but to investigate whether different sets of rate parameter values could describe the same mechanical behavior and whether parameter values could be identified that accurately predict both mechanical and metabolic behavior. Furthermore, the magnitude of the predicted CE-contraction velocities was around 0.3 *lce*_*opt*_ /*s*, which was low relative to the maximum shortening velocity. This was also the case for van der Zee et al. (2019), who estimated CE-contraction velocities to be around 0.3 *lce*_*opt*_/*s* as well. We do not expect a substantial number of mechanical detachments to occur at such low contraction velocities. Thus, we expect the potential effect of mechanical detachments on the predicted metabolic power in this study to be minimal.

The low contraction velocities are also relevant for the other simplification that we made: the exclusion of metabolic power needed for muscle activation. During the dynamic condition, the muscle activation increases during shortening and decreases during lengthening to maintain a constant torque. However, the CE force-velocity relationship is non-linear (Hill, 1938), thus the magnitude of the increase in activation during shortening will generally not be equal to the magnitude of the decrease during lengthening. Only a small part of the force-velocity relationship around the isometric point, a part which was approximately linear, was visited during the simulations. Thus, the magnitude of the increase in activation during shortening and the decrease during lengthening were similar. This resulted in the normalized calcium concentration, averaged over time and the 10 values of *g*_1_, to be similar during the dynamic condition and the isometric condition (0.102 for the dynamic condition and 0.100 for the isometric condition, see Fig. 2(A)). The predicted metabolic power needed for muscle activation would therefore have been similar between the two conditions if it was included in the model. Thus, the inclusion of the metabolic power needed for muscle activation would not have substantially influenced our results.

### Formulation of the rate functions

The same mechanical behavior cannot only be described using different sets of rate parameter values for a given formulation of the rate functions, but also by using different formulations of the rate functions (Huxley, 1957; Lemaire et al., 2025). In this study, we chose a formulation that was computationally simple, but we could have used other formulations that allow for equally good descriptions of the steady-state force velocity relationship (albeit at different rate parameter values). However, choosing other rate function formulations would not have changed our findings. After all, the key observation is that for many viable formulations of the rate functions, there will also be multiple sets of rate parameter values that will lead to accurate descriptions of the mechanical behavior. Although the formulations of the rate functions used in this study were sufficient for the experimental conditions simulated here, they may not be adequate for other experimental conditions where both the mechanical and metabolic behavior are of interest. Thus, we advise researchers that aim to use Huxley-type models for predictions of mechanical and metabolic muscular behavior to keep this in mind while choosing the shape of the rate functions.

### Implications for metabolic power predictions using Huxley-type muscle models

When the goal is to accurately predict both mechanical and metabolic behavior of muscles, Huxley-type muscle models might be preferred over Hill-type models as Huxley-type models contain a conceptual link between mechanical and metabolic behavior in the form of cross-bridge cycling. When using Huxley-type models to describe an *in vivo* experiment, it might be tempting to only fit the cross-bridge rate parameter values to data on the mechanical behavior and then assume that predictions of the metabolic behavior will also be accurate due to the aforementioned link between mechanical and metabolic behavior in the model. However, our results show that this assumption is incorrect, as different sets of parameter values can lead to nearly identical mechanical behavior while yielding substantially different metabolic behavior. Instead, the existence of multiple rate parameter sets that produce equivalent mechanical behavior can be exploited to simultaneously optimize both mechanical and metabolic behavior.

In Fig. 4, we saw that the number of detachments at *x*/*h* < 0 and *x*/*h* > 1 was slightly larger and, more importantly, that the number of detachments was lower between *x*/*h* = 0 and *x*/ *h* = 1 when *g*_1_ = 20 Hz compared to when *g*_1_ = 100 Hz. These two effects of decreasing *g*_1_ explain why the predicted relative difference in metabolic power between the two conditions increased with a decreasing value of *g*_1_ (Fig. 4). Importantly, the finding that the best match of the predicted and experimentally measured difference in metabolic power occurred at a low value of *g*_1_ (37 Hz) suggests that the number of cross-bridge detachments during isometric contractions is relatively low. Furthermore, the predicted number of detachments between *x*/*h* = 0 and *x*/*h* = 1 was almost equal between the isometric and dynamic conditions. Interestingly, these findings thus suggest that the large increase in metabolic power measured experimentally in the dynamic condition would be almost exclusively due to the additional detachments at *x*/*h* < 0 and *x*/*h* > 1.

For future studies investigating if a set of rate parameter values can be found that leads to accurate predictions of mechanical and metabolic behavior across a wide range of conditions, it would be good to use experimental conditions in which the primary difference between the conditions is the average contraction velocity (e.g. Swinnen et al. (2023)). It is well known that contraction velocity has an effect on metabolic power consumption (Fenn, 1924; Hill, 1938) and that, in terms of the Huxley-type model, the increase in metabolic power consumption with increasing shortening velocity can be explained by an increase in the number of cross-bridge detachments (Huxley, 1957). In this study, we considered two distinct conditions of a specific task, and we found that for this set of conditions we can find a set of parameter values that describes both the metabolic and mechanical behavior well. However, it remains to be seen whether similar results can be obtained for a broader set of movement conditions.

## Conclusion

Using a Huxley-type muscle model, we showed that different sets of cross-bridge rate parameter values will lead to different predictions of metabolic behavior. At the same time, we showed that the same mechanical behavior can be described accurately using sets of cross-bridge rate parameter values in which the imposed (*g*_1_) varied 10-fold. This flexibility can be exploited to obtain accurate predictions of mechanical and metabolic behavior simultaneously. It would be interesting to further explore to what extent a single set of rate function shapes and corresponding rate parameter values can describe currently available data on mechanical and metabolic behavior in a wide range of conditions using a Huxley-type muscle model.

## Authors contributions

Roeland T. Vos: Conceptualization, Methodology, Software, Validation, Formal analysis, Investigation, Data curation, Visualization, Writing – original draft.

Koen K. Lemaire: Conceptualization, Methodology, Supervision, Funding acquisition, Writing – review & editing.

Luuk Vos: Conceptualization, Methodology, Writing – review & editing.

Arthur J. van Soest: Conceptualization, Methodology, Supervision, Writing – review & editing.

Dinant A. Kistemaker: Conceptualization, Methodology, Supervision, Writing – review & editing.

## Data, scripts, code and supplementary information availability

All MATLAB code and data required to reproduce the analyses and simulations presented in this manuscript are available in the repository at DOI: https://doi.org/10.5281/zenodo.21427753.

